# A Collective exponential attractor shapes growth-rate variability in single bacterial cells

**DOI:** 10.1101/2024.06.23.600237

**Authors:** Kuheli Biswas, Amy E. Sanderson, Hanna Salman, Naama Brenner

## Abstract

Exponential accumulation of cell size and highly expressed proteins is observed at the single-cell level in many bacterial species. While the exponential rates fluctuate from cycle to cycle, they remain stable on average over time and strongly correlated across different proteins and cell size. In this study, we investigate growth-rate variability at this state of *balanced biosynthesis*, and present a theoretical framework to explain its properties through the emergence of a high-dimensional collective dynamic attractor. This stable attractor arises from recurrent interactions among multiple cellular components, driving them to converge to the same exponential growth rate, thereby sustaining the balanced state. The convergence of growth rates induces a decay in instantaneous growth rate noise throughout the cell cycle, with a faster decay for higher average growth rates. Notably, our analysis identifies random deviations from symmetric division as the primary source of growth rate variability. The theory offers a coherent set of predictions for many observations, validated by extensive experimental single-cell data. The spontaneous emergence of homeostasis through dynamic interactions suggests that specific control mechanisms to compensate deviations from a target may not be necessary to maintain homeostasis in a balanced state.

## Introduction

Exponential growth is a hallmark of proliferating bacterial populations (1). It results from cell divisions occurring repeatedly, at a constant average rate, which in turn cause a doubling of the population at fixed time intervals. At balanced exponential growth, all extensive properties such as biomass and cell number accumulate exponentially at the same rate (2–4). Notably, this population-level behavior does not necessarily require individual cells to grow exponentially. Indeed, while exponential growth at the population level was studied for decades, the quantitative nature of individual cell growth remained unclear. Single-cell measurements have since revealed that many types of bacteria and eukaryotic cells accumulate mass in a supra-linear manner, well-describe by a single exponent (5, 6).

Exponential growth at the single-cell level is difficult to distinguish from other growth patterns due to the small dynamic range -approximately twofold over a cycle. Some experiments have detected measurable deviations from a single exponent, reflected as changes in instantaneous growth rate across the cell cycle (7–10). Still, a large body of data across diverse cell types exhibits growth patterns for which a single exponent remains an excellent approximation, defining an effective variable for the cycle (6, 9, 11–19). This effective growth rate can fluctuate considerably from one cycle to the next, but remains stable on average across many cell divisions in a lineage (6). Highly expressed proteins exhibit a similar behavior: their accumulation during each cycle is approximately exponential, with a rate that fluctuates across cycles but is stable on average. These properties are crucial in shaping the universal steady-state distributions of cell size and of protein content across lineages (20–22), as well as the non-universal distribution of times between divisions (23).

Importantly, examining both cell size and protein levels within the same cell reveals a single-cell version of balanced exponential growth. The exponential rates of all these different phenotypic components are strongly correlated on a cycle-by-cycle basis (11, 16, 18). This observation, often termed balanced biosynthesis (4), could result from control mechanisms that coordinate the different components. Another possibility is that exponential growth rate is a global property of the cell as a whole, emerging collectively from recurrent interactions among its multiple components.

Why is exponential accumulation so prevalent in growing cells of different types? What are the microscopic events that govern the fluctuations in growth rates across cycles in a lineage? Can a single dominant source of these fluctuations be identified? and how are the exponential rates of different cellular components coordinated? In this work, we analyze statistical properties of growth rates - both effective across an entire cycle, and instantaneous - to shed light on these questions. We propose that these statistical properties contain information that can distinguish between different potential answers. Combined with recent theoretical results of multi-dimensional cellular models, we present a unified physical picture where exponential growth is an emergent property induced by an exponential attractor of the collective dynamics in a high-dimensional phase space. In this picture, fluctuations in growth rate are dominantly influenced by imperfect cell division events, which perturb the growth trajectory off the attractor in random directions. This framework explains growth and division homeostasis in a purely dynamic manner, without the need to rely on engineered feedback control. It offers several specific predictions that are tested and borne out in single-cell data.

### Empirical properties of growth rates

To set the stage for our analysis, we first review several established statistical properties of single-cell growth rates, as well as their precise definitions. Single-cell experiments, which track growing and dividing cells for multiple generations, have revealed that biomass and highly expressed proteins accumulate approximately exponentially between divisions. An example is shown in Fig. 1a and b respectively, depicting *E. coli* ‘s growth trajectories in a mother-machine microfluidic device (data from (16)). Exponential fits to both cell size and protein accumulation are in good agreement with the data, defining an effective growth rate (eGR, *λ*) over each cell cycle. These eGRs vary from one cycle to the next, with strong cycle-by-cycle co-variation between components. Such co-variation was found between cell size and proteins, and between two proteins measured in the same cell, for arbitrary tagged proteins (9, 11, 16, 18). An example is depicted in Fig. 1c; we refer to this single-cell property as “balanced biosynthesis”.

**FIG. 1.**
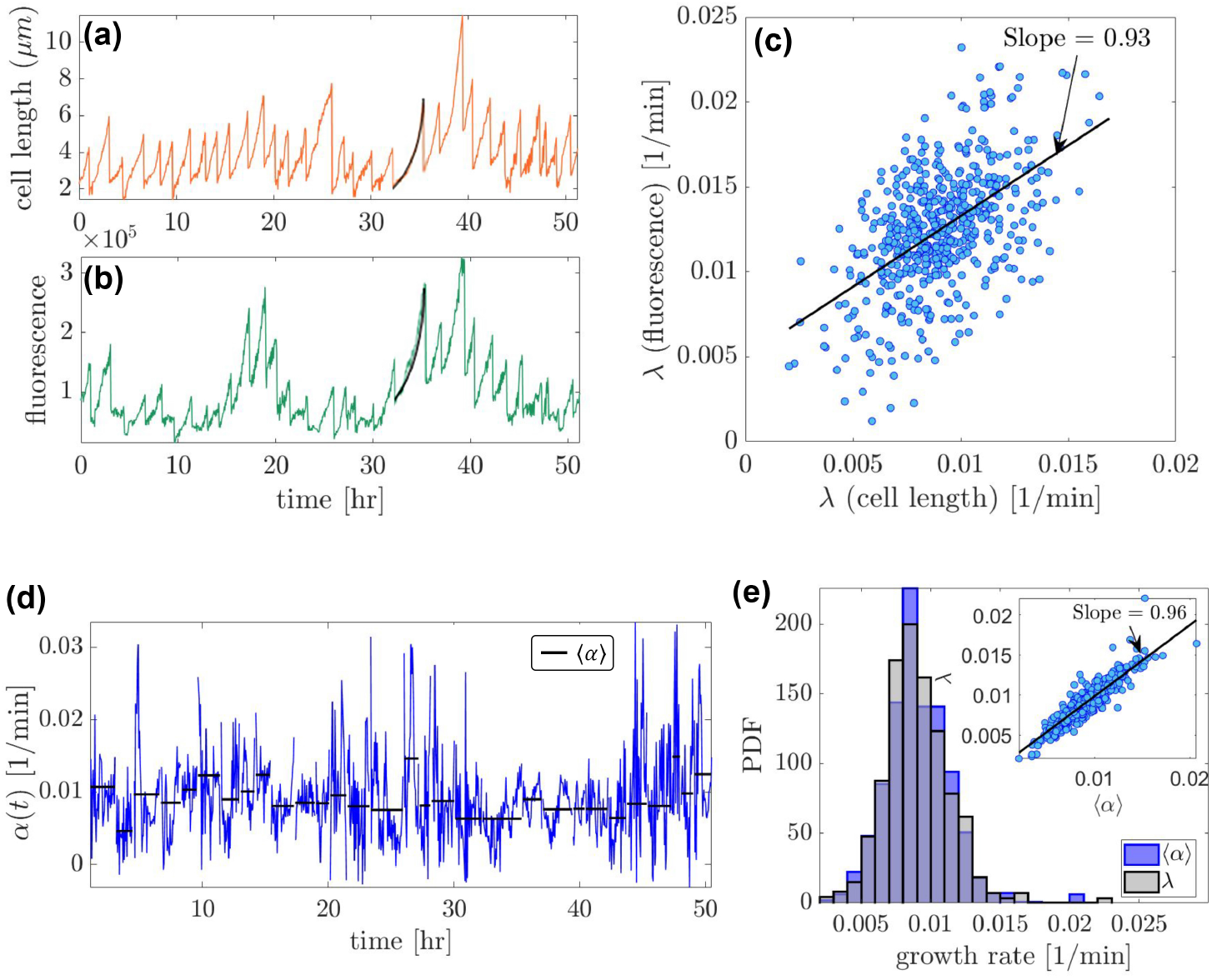
Definitions and empirical properties of growth rates. Time traces of cell size **(a)** and protein content **(b)** in *E. coli* cells growing in a mother-machine microfluidic channel. Effective growth rates (eRG), *λ*, are defined by fitting the dynamics during each cell cycle with exponential functions (black lines). **(c)** *λ*s of cell size and of protein copy number are correlated across cycles, here with coefficient 0.64. **(d)** The instantaneous growth rate (iGR) is the logarithmic derivative of the continuous signal, here estimated for cell size across time (blue line). One may estimate also the average of iGR over each cycle, ⟨*α*⟩ (black steps). **(e)** eGR *λ* and averaged iGR ⟨*α*⟩ have similar distributions, and are strongly correlated (inset).

The distribution of cell size eGR, collected across cycles and/or across lineages, has been reported approximately Gaussian-shaped. The level of variability, quantified by the coefficient of variation (CV), changes significantly among published results, ranging from 5% to 40% (12, 13, 16– 19, 24). Our recent study found that this broad range of eGR variability markedly affects the division time distribution, accounting for its non-universal shape across experiments (23). At the same time, the primary factors underlying eGR variability itself still remain unclear. Modeling work has suggested several possible origins of these fluctuations, including the intrinsic molecular noise resulting from the synthesis and degradation of cellular components and random distribution of mRNAs, and random deviations from symmetric division (25, 26). Identifying a dominant source of eGR variability has proven challenging. In particular, the large range of CV seen in different experiments, some of which were performed in similar conditions, remains puzzling.

Complementary to the discrete eGR defined over a cell cycle, one may characterize single-cell growth by an instantaneous rate - iGR - the logarithmic derivative of continuously measured cell size or protein or any other component. For purely exponential and deterministic growth, the iGR is constant over time and equal to the eGR. In experimental data, where both deviations from a perfect exponential and noise exist, iGR fluctuates in time as seen in Fig. 1d. One may define the time-averaged iGR over a cycle duration, marked by black steps in Fig. 1d; this quantity is tightly correlated with the eGR obtained by fitting. Fig. 1e shows that the distributions of these two quantities across cycles is similar, and that their cycle-by-cycle correlation is high (0.87; inset).

### Models of balanced exponential growth

What is the source of coordinated exponential accumulation in cellular components and size? Exponential growth is associated with autocatalytic processes, such as ribosome proliferation inside the cell. However, it can also arise indirectly by multi-dimensional recurrent interactions. Linear interactions generally lead, at long times, to exponential growth of all components at a rate governed by the dominant positive eigenvalue of the interaction matrix (16, 27, 28). Recent theoretic work has shown that a similar property is shared by a broad class of models with *nonlinear* interactions (25, 29). Included in this class are all mass-action kinetics, making it very general and relevant to bacterial cells. Cell size is assumed to be a linear combination of its components, thus growing with the same exponential rate, and inducing a stabilizing negative feedback on the concentrations which in turn govern the reaction rates. Thus, in this class of models balanced biosynthesis is a property that emerges from the collective dynamics and no control mechanisms to coordinate growth rates is required (see Supplementary Note 1).

Several specific models in this class were studied previously (18, 25, 26, 30–32). Here we use a special case example (following (25)) to demonstrate the properties of the entire class of models. The example consists of a coarse-grained description of three cellular components (Fig. 2a): amino acids, metabolic enzymes, and ribosomes with copy numbers 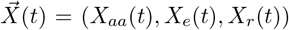.The cell size is given by their combination *V* (*t*) = *ρ*_*aa*_*X*_*aa*_(*t*) +*ρ*_*e*_*X*_*e*_(*t*) +*ρ*_*r*_*X*_*r*_.(*t*), so that concentrations are 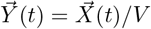.Interactions among components are governed by the dynamical system

**FIG. 2.**
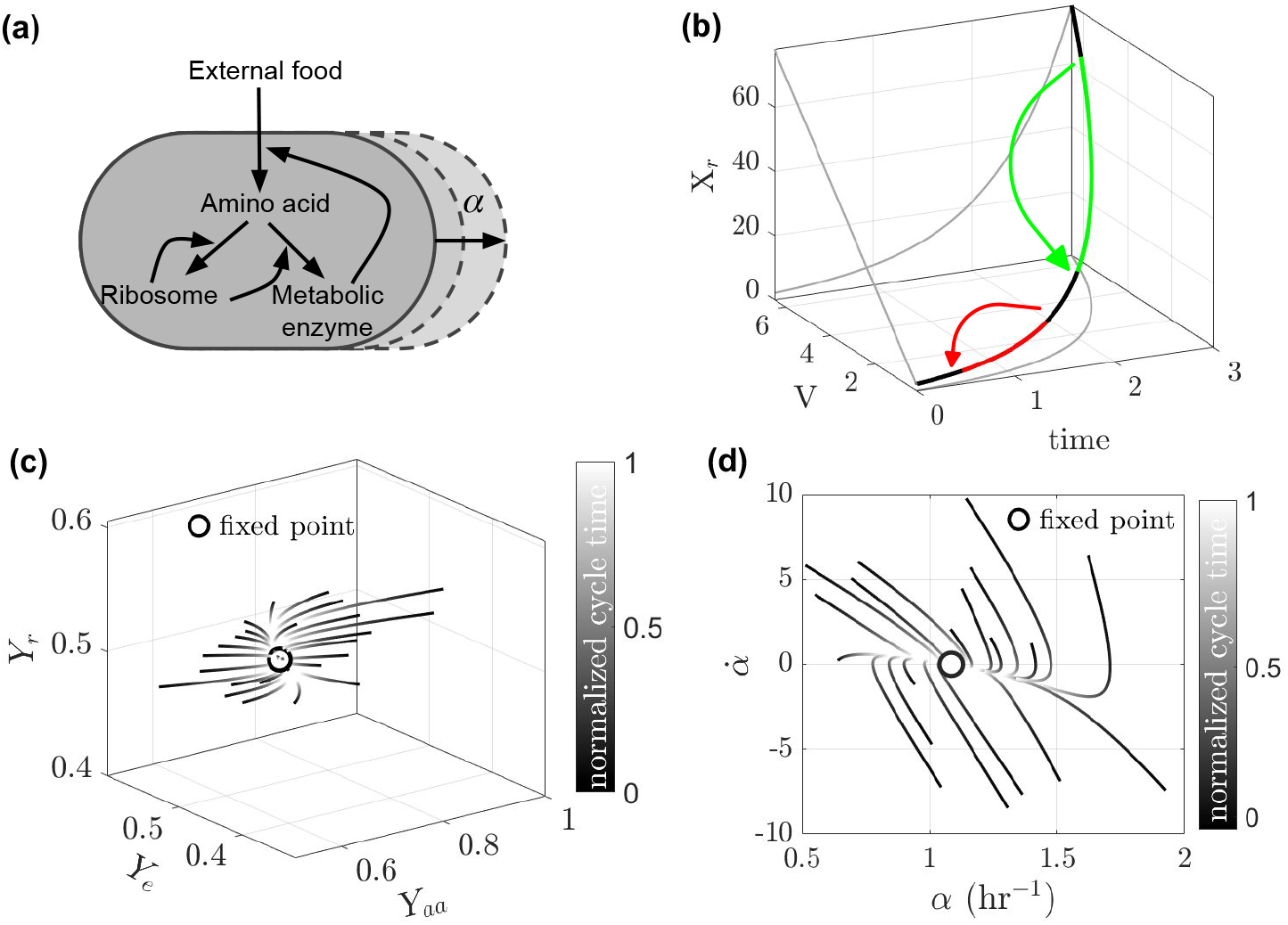
Collective exponential attractor from multi-dimensional nonlinear interactions. **(a)** A three dimensional kinetic model represents a coarse-grained cell (Eqs. 1). It is a specail case of a large class of models with similar properties. **(b)** The trajectory of the resulting dynamics from the model simulation is presented as a black line in the three-dimensional projection space including one of the components (*X*_*r*_), the system size - volume (*V*) and time. All components maintain a fixed ratio to the volume, reflected by the straight line in the (*X*_*r*_, *V*) plane; any two components also maintain fixed ratio, both during growth and after symmetric division. Two cycles of continuous growth on the collective attractor (color on the black attractor) and exactly symmetric division (arrows), are shown in red and green for two initial conditions. **(c)** The collective exponential attractor maps onto a stable fixed point in concentration space. Gray lines: trajectories initiated at random initial conditions. circle: fixed point of the dynamics. **(d)** Phase plane of instantaneous growth rate 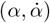, illustrating the convergence to the fixed point in term of growth rate dynamics. Also here trajectories are initiated at random points in the plane. In **(c)** and **(d)** time is encoded by gray-scale. For parameter values of the simulation see Supplementary Table 1.

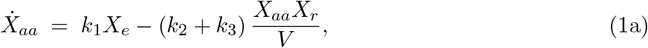

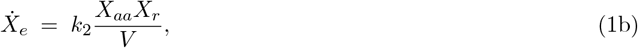

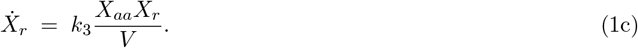

In this reaction network, amino acids are produced by the catalytic action of the metabolic enzyme from external food molecules at a rate *k*_1_. Enzymes and ribosomal proteins are synthesized from amino acids in reactions catalyzed by ribosomes with rate *k*_2_ and *k*_3_, respectively. This model obeys a scalability condition (25, 29), ensuring that at long times the dynamics are attracted to a trajectory where all components accumulate exponentially at the same rate *α*. We show in Fig. 2b the projection of this concerted exponential growth (black line) on the sub-space of one cellular components, cell size and time, (*X*_*r*_, *V, t*). The exponential accumulation of ribosomes, for example, is apparent on the projection plane (*X*_*r*_, *t*). Since all components and cell size grow with the same rate, ratios between any two components is conserved, as well as ratios between any component and cell size *V*. This property is reflected on the projection (*X*_*r*_, *V*) in Fig. 2b, where a straight line appears *V* = *cX*_*r*_.

Symmetric division of all components respects the ratios between them, and following such a division event the trajectory jumps to another point on the same attractor, where all components are half of their values prior to division. Therefore, repeated cycles of continuous growth and discrete divisions can take place without ever leaving the exponential attractor. These properties remain true regardless of the mechanism or criterion for division. Importantly, they remain true for any initial condition on the attractor, all having the same ratios between components but different absolute values. Two examples of such cycles are depicted in Fig. 2b in red and green, highlighting this degeneracy of the cycles. Cell size, being a combination of these components (*V* = Σ_*i*_*ρ*_*i*_*X*_*i*_), also varies between degenerate solutions on the attractor. This observation can explain the significant variability in time-averaged cell size across distinct lineages in similar conditions (16, 24).

A key property of the collective exponential attractor described above is its stability (25, 29): not only do trajectories that start on it remain on it, but following deviation they are drawn back to it over time. This stability is particularly evident in the space of concentrations, *Y*_*i*_ = *X*_*i*_/*V*, where the exponential attractor is mapped onto a single fixed point (29), acting as a sink for nearby trajectories. Figure 2c illustrates this for our test-case model, in three-dimensional concentration space, where trajectories initialized in different locations in its vicinity (gray lines, with gray-scale indicating time) relax back to the fixed point (marked by a circle).

The exponential growth rate 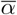 is common to all components on the attractor. Therefore, convergence to the stable fixed point will be accompanied by convergence of the instantaneous growth rate *α*(*t*) to its fixed point value. Fig. 2d shows simulated trajectories of the dynamics in the phase plane 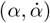, illustrating the convergence of growth rate to its fixed point. For our example, the explicit fixed point and the corresponding eigenvalues and eigenvectors are computed in Supplementary Note 2.

Importantly, all the features described above and illustrated for a special case, are shared by a large class of models relevant to cellular dynamics (25, 29). In what follows, we rely on these features to formulate quantitative predictions for statistical properties measured in single-cell experiments, and compare them to data. The generality of the predictions, and their insensitivity to model details, make the comparison especially meaningful.

### Model predictions and comparison to data

A key aspect of comparing our model to data is understanding how cell division influences growth dynamics. Recall that the attractor is invariant to perfect division which preserves ratios; in contrast, noise in division causes imperfect ratios between components that push the trajectory off the attractor. Due to its intrinsic stability, coordination is re-established in the next growth cycle, pulling the trajectory back. Over multiple cycles, imperfect division repeatedly perturbs the system and spreads the initial conditions of individual cycles, which then converge back toward the attractor. Figures 2c and d were generated from dynamic simulations incorporating noisy division, where division occurs following a threshold-crossing trigger, and daughter cells receive a fraction *f* of each component, drawn from a Gaussian distribution centered at 1/2 with CV_*f*_ = 15% (9, 12, 13, 17–19, 24). Regardless of implementation details (such as division noise level or exact division condition), as long as division randomly spreads initial conditions, consecutive cycles exhibit the characteristic convergence to the fixed point (Supplementary Note 4).

One immediate consequence of this dynamic picture, is that individual cell cycles start out very different from one another and gradually become more similar, as the cycle progresses and they are all drawn back to the same attractor. This is expected for both concentrations and iGRs, which are properties uniquely defined on the exponential attractor, but not for absolute quantities for which the attractor is degenerate, such as cell size or amounts of different components. To quantify this effect, we estimate the Coefficient of Variation (CV, (standard deviation)/mean) across individual cycles. Such an analysis has revealed a decrease of growth rate variability in *B. subtilis* (9). Similar behavior for *E. coli* across several different experimental conditions is presented in Fig. 3a. To compare different data sets, the cell cycle was normalized by its mean for each experimental condition; in all cases a marked decrease of CV_*α*_(*t*) is observed. To further test this prediction, we followed filamenting *E. coli* cells in the mother machine, expecting to see a further decrease of CV_*α*_(*t*) as the dynamic range is extended and the iGRs continue to approach the growth attractor without disruption by divisions. This prediction is observed clearly in Fig. 3a. Analyzing protein concentration in the same way, Fig. 3b shows that *CV*_*Y*_ also decreases as the cycle progresses. Importantly, the same analysis for cell size does not show such a decrease (Supplementary Fig. 1). Another prediction that can be derived from the model concerns the rate at which CV_*α*_(*t*) decreases along the normalized cycle time: this decrease is faster in faster-growing cells compared to their slower-growing counterparts. This is demonstrated by data from *E. coli* cells grown at different temperatures (12) and in different nutrient conditions (18) (Fig. 3c), in agreement with model calculations ((Fig. 3c, inset). Quantifying the decay rate in iGR noise by fitting *CV*_*α*_(*t*) to a decreasing exponential function, we examined the resulting decay rate as a function of the corresponding mean growth rate across a host of experimental data. This analysis reveals an increasing dependence, displayed in Fig. 3d, as predicted by the model (inset).

**FIG. 3.**
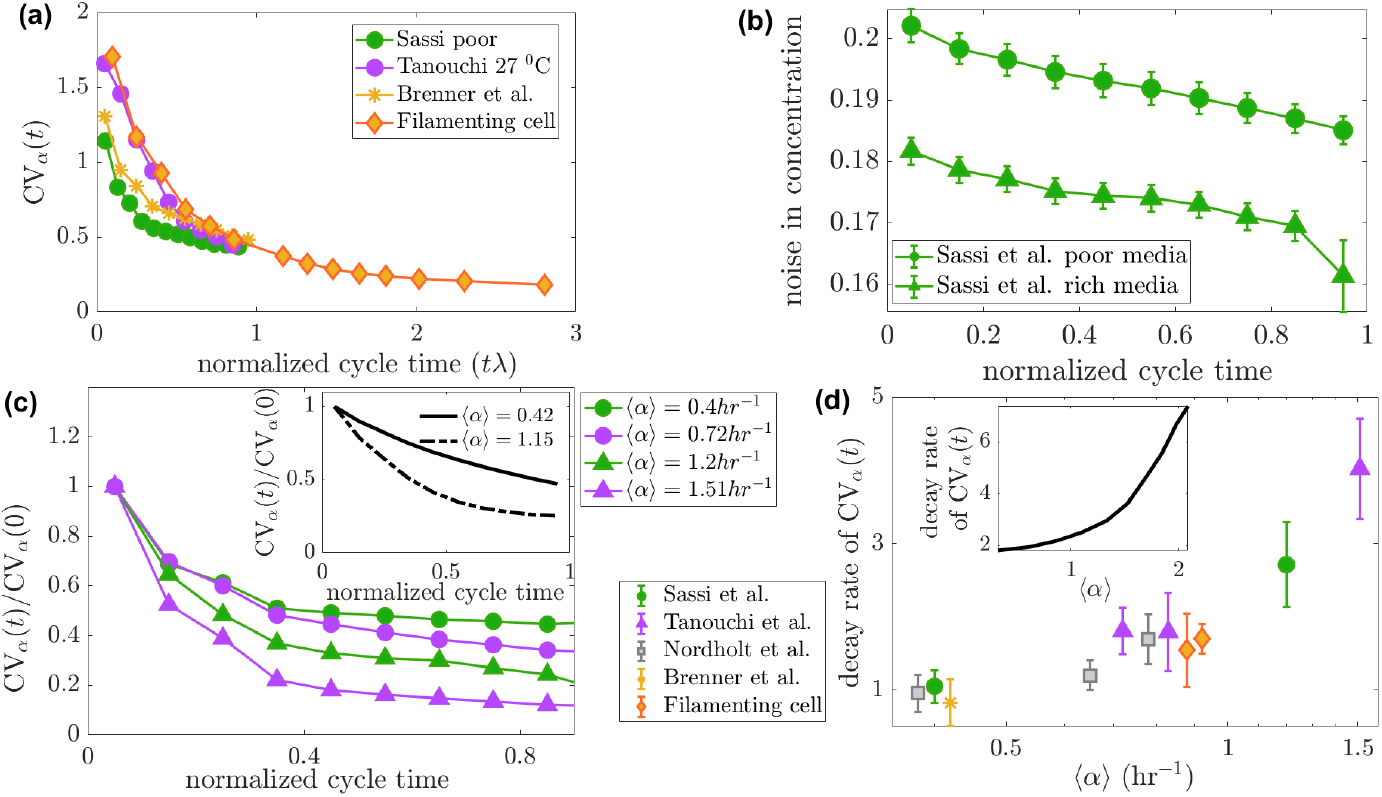
Testing Model Predictions in Data. **(a)** iGR noise across individual cycles (CV_*α*_) decreases along the cycle duration, both for wild-type (green (18), violet (12), yellow (11)) and for filamenting cells (orange diamonds). Time is normalized by mean growth rate. CV_*α*_ is estimated by segmenting the data into bins containing equal numbers data points, calculating the standard deviation and mean iGR within each bin, and computing their ratio. **(b)** *CV*_*Y*_ for protein concentration along normalized cell cycle time. **(c**,**d)** iGR noise decreases more rapidly for faster growing cells. Shown in **(c)** are plots of CV_*α*_ normalized by their initial value, from *E. coli* data measured under different nutrient conditions (18) and temperatures resulting in different mean growth rates. **(d)** displays the decay rate, best fit exponential to *CV*_*α*_(*t*) in plots such as those in **(c)**, with confidence intervals (error bars) as a function of mean growth rate in each condition. Experimental details on filamenting cells are given in Supplementary Note 3, and on dividing cells in Supplementary Table 2

From the data presented thus far, we conclude that division noise can explain several features of iGR statistics, as predicted by our model where division is the only noise source. A more nuanced question is whether we can assess the importance of division noise relative to other potential noise sources. To this end, we added fluctuations in all components to the model, which represent intrinsic molecular noise and in turn induce instantaneous fluctuations in iGR. This allows us to modulate the relative strength of the two stochastic effects, division and molecular, and asses their contribution to the measurable statistical properties of iGR and eGR. Figure 4 shows the resulting dynamics of iGR in two model simulations, where either intrinsic molecular noise **(a)** or division noise **(b)** is dominant. Estimating the variability in iGR similar to the analysis in Fig. 3, shows that the decrease in *CV*_*α*_, characteristic of the data, appears only in the regime of dominant division noise (Figure 4c). This indicates that the data are in this regime, and that division noise plays a crucial role in the observed statistical properties.

**FIG. 4.**
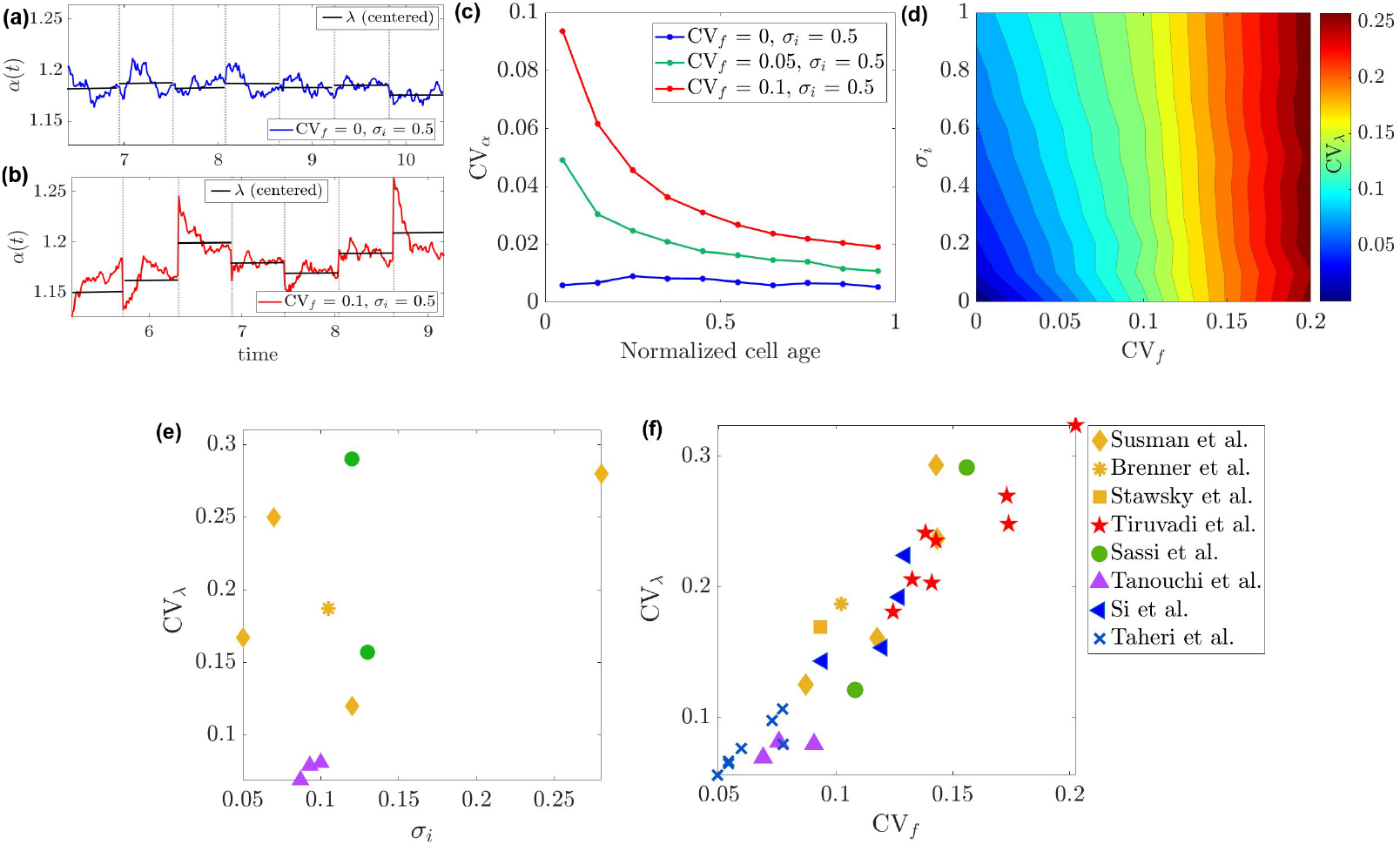
Contribution of different noise sources to growth-rate variability. **(a,b)** Model simulations of iGR dynamics in two regimes, where molecular noise **(a)** or division noise **(b)** is dominant. **(c)** Model simulations of iGR noise over normalized cell cycle for different parameter values (legend). *σ*_*i*_ is the level of intrinsic molecular noise and CV_*f*_ is division noise. The decrease in CV_*α*_, observed in experiment, appears only in the region *σ*_*i*_ *<<*CV_*f*_. **(d)** Model calculation of eGR varibility as a function of the strength of two noise sources. **(e**,**f)** eGR variability estimated from data as a function of intrinsic molecular fluctuation strength and of division noise strength. The intrinsic molecular noise is estimated from the data as *σ*_*i*_ = std[*V* (*t*) − *V*_fit_(*t*)], where *V* is the cell volume. For the model, noise is added to the dynamic equations, 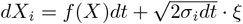,where *ξ* is standard normal Gaussian noise.

We next consider statistical properties of the discrete eGR, defined over entire cycles. Since division noise perturbs the dynamics on the exponential attractor, the growth trajectory during each cell cycle is not an exact exponential any more. However, due to the small dynamic range (twofold in all components), it can still be approximated by a single exponential rate; these effective rates fluctuate from one cycle to the next. The property of balanced biosynthesis now becomes statistical in nature - accumulation rates are not identical but are highly correlated, faithfully matching experimental results ((11, 16, 18); see also Fig. 1). This theoretical picture implies that as noise in division increases, eGR variability should becomes larger. To estimate the relative importance of the two noise sources on eGR, we used the model to compute *CV*_*λ*_ in the parameter plane of their strengths. Fig. 4d shows that this measure of eGR variability depends strongly on division noise *CV*_*f*_ but weakly on intrinsic molecular noise *σ*_*i*_.

To test this prediction with respect to experiment, we assembled a broad array of single-cell measurements and plotted the empirically estimated eGR noise - *CV*_*λ*_ - as a function of instantaneous fluctuations *CV*_*α*_, and of division noise, *CV*_*f*_. The results are displayed in Fig. 4d,e, showing a stronger correlation with division noise than with intrinsic molecular noise. To consider other possible noise sources that could contribute to the observed variability in effective growth rates, we incorporated in our model stochasticity in the threshold triggering division, as well as intrinsic number fluctuations in chemical reactions. Notably, we find the impact of threshold noise on top of division noise is negligible (in agreement with (25)). Additionally, simulating the discrete version of the chemical reactions to account for the effect of a small number of molecules, shows that number fluctuations become important for eGR variability only in the region where division noise is very small; in the presence of realistic division noise, their additional effect is insignificant. These results are presented in Supplementary Fig. 2. Comparing this to Fig. 4f, we find that the sharp decrease of eGR variability at small division noise indicates that number fluctuations do not play a central role here. This is consistent with our model that describes coarse-grained cellular dynamics by highly abundant components which accumulate smoothly over the cycle; in such variable, number fluctuations are expected to be negligible.

### Summary and Discussion

The mathematical foundations for understanding sustained exponential growth in natural systems, such as cells with nonlinear reaction networks, have recently been clarified (25, 29). A broad class of models was identified that demonstrates asymptotic coordinated exponential growth for all components, with cell size growing at the same rate as a weighted sum of these components. These features appear with weak sensitivity to parameters or model details, and rely on a negative feedback loop between cell size and its constituents that ensures system stability: accumulation of components activates cell growth, but their buildup is inhibited by dilution as cell size increases. This give a robust theoretical framework to study multi-dimensional exponential growth in cells and their constituents.

In this work, we investigated the statistical properties of single-cell growth rates in light of this theoretical framework and the emergent collective exponential attractor. Using a specific set of equations as a representative of the class of models, we investigated the interplay between the deterministic system’s globally stable exponential attractor and the stochastic division events. Division noise perturbs the deterministic growth trajectories away from the attractor, while continuous growth during the cell cycle relaxes the system back to equilibrium.

Empirically, growth rate can be measured either as the continuous logarithmic derivative of protein or cell size (iGR), or through an effective fit over entire cell cycles (eGR). While strongly related, these two quantities offer different insights into the system’s dynamics. Both can be faithfully estimated from single-cell measurements of cells across time as they grow and divide. The collective exponential attractor shapes the statistical properties of both these phenotypes; the abundance of available data, including different cell types and conditions, allows for a meaningful testing of the theoretical framework with experiment.

We identified three key predictions of the theory. First, the coefficient of variation (CV_*α*_) of instantaneous growth rate (iGR) across individual cell cycles decreases over the cycle, a phenomenon previously observed for *B. subtilis* (9) and now shown to be more general and explained through the attractor framework. Second, iGR variation decreases more rapidly in faster-growing cells, as relaxation rate scales with the mean growth rate. These predictions are validated by empirical data from various experimental studies on bacterial cells, including our data on filamenting *E. coli*, and align with previous work (16, 25). We further investigated the relative contributions of division noise and intrinsic molecular fluctuations to eGR variability, leading to the third prediction: division noise is the primary contributor, dominating over intrinsic noise as well as other potential noise sources. This may be a surprising result since division is often considered symmetric with high precision in *E. coli* ; it is nevertheless supported by the experimental data, as well.

Taken together, our results suggest that the single-cell biomass (or volume) growth rate is a global phenotype shaped by the collective dynamics of many components. As an alternative, one may imagine a situation where growth rate is limited by a single or few molecular components - a growth-rate determining molecule. In the class of models we consider, the iGR of cell size depends on changes in their concentrations. At the asymptotic attractor, where concentrations stabilize, the growth rate is shared by all components and is determined by system parameters. In the presence of noise, this leads to growth rates of different components being correlated but not identical.

This view is supported by previous work, although often stated in different contexts. The recurrent relationship between cell growth fluctuations and protein expression noise was highlighted in (33). While a different theoretical model was developed in that study, its qualitative conclusion was that the entanglement between instantaneous growth and expression noise reflects the inherent autocatalytic nature of a self-replicating system, in line with our view. More recent work highlighted the central role of cell division as a disruption to spatially coordinated cellular dynamics (34). Also there the physical effect and model are different, but the basic idea is the same: small deviations from symmetric division have consequences on growth rate that extend beyond their quantitative noise level, when viewing the cell as a multi-dimensional system. These deviations can disrupt collective dynamics, be it through ratios or through spatial cellular organization, that require a re-buildup following division.

The recovery from the system-level disruption by division has been viewed as evidence for dedicated compensation mechanisms underlying homeostatic regulation (9, 34). Our results suggest that such recovery and the resulting long-term stability can emerge from stable high-dimensional dynamics of interacting components. Negative multi-level global feedback, such as appears in the dynamic models we study, ensures homeostasis across division cycles, offering an alternative to control theory where local estimation and compensation are required. This systems-level dynamic homeostasis does not depend on fine-tuned interactions or parameters but on gross properties of the class of models. Indeed, the results we discuss are independent of the exact coarse-grained interactions contained in the model, which regulate cellular growth, or the specific mode of cell division and its regulation. The questions of which components and what precise interactions determine cellular growth and regulate cell division, have been the topic of much study and debate and further research is still needed to address them. However, our approach allows to separate this problem from the understanding the coarse-grained phenomenology of stable exponential growth correlated among the different cellular characteristics and resulting statistical properties.

