## supplementary text and figures for "A Collective exponential attractor shapes growth-rate variability in single bacterial cells"


### SUPPPLEMENTARY NOTES

#### Supplementary Note 1: General Model Formulation

Recent theoretical studies have shown that exponential growth with a single rate emerges in a broad class of models describing systems of non-linearly interacting components, such as chemical reaction networks (1, 2). For completeness we summarize here some of the results. Imagine the cell as a container of volume $V(t)$ containing $n$ components, with copy numbers $X_i,\ i = 1, 2...n$. Dynamics of the $i^{th}$ component is given by

$$\dot{X}_i = F_i(\mathbf{X}), \tag{1}$$

where $F_i$ are nonlinear functions, representing the system's chemical reactions. The cell size is a linear combination of its components, $V(\mathbf{X}) = \sum_{c=1}^{n} \rho_c X_c$. In such a system, one finds the emergence of an asymptotic *exponential attractor* under the following condition on the functions $F_i(\mathbf{X})$:

$$F_i(\beta \mathbf{X}) = \beta F_i(\mathbf{X}) \quad \text{for any } \beta > 0. \tag{2}$$

The condition is obeyed by functions that describe mass-action kinetics of various order, and thus is very relevant to models of cellular interactions. On the exponential attractor all components and cell size grow exponentially at the same rate after a very long time (asymptotically). Consider two components $X_i, X_j$ of the trajectory on the attractor: if their ratio at time $t = 0$ is given, then that ratio is maintained for all times

$$\frac{X_i(t)}{X_j(t)} = \frac{X_i(0)e^{\overline{\alpha}t}}{X_j(0)e^{\overline{\alpha}t}} = \frac{X_i(0)}{X_j(0)}. \tag{3}$$


[*]

[†]

Where $\overline{\alpha}$ is the attractor growth rate. Additionally, the concentrations $Y_i = X_i/V = Y_i/\sum_{k=1}^{n} \rho_k X_k$ are also conserved for all $i$. To analyze this dynamical system in the space of concentrations instead of copy numbers, we write Eq. (1) in terms of concentrations, including also the effect of dilution by expanding volume:

$$\dot{Y}_i = \frac{1}{V} F_i(V\mathbf{Y}) - \alpha(t)\, Y_i, \tag{4}$$

where the volume growth rate, is defined as

$$\alpha(t) = \frac{\dot{V}}{V} = \sum_{i=1}^{n} \rho_i F_i(\mathbf{Y(t)}). \tag{5}$$

#### Supplementary Note 2: Explicit calculations for our special-case model

In our work we consider a concrete example involving a non-linear interaction network of three components: amino acids, metabolic enzymes, and ribosomes with respective population size $X_{aa}$, $X_e$, and $X_r$. The equations governing the system are

$$\begin{aligned} \dot{X}_{aa} &= k_1 X_e - (k_2 + k_3)\,\frac{X_{aa} X_r}{V}, & \text{(6a)} \\ \dot{X}_e &= k_2 \frac{X_{aa} X_r}{V}, & \text{(6b)} \\ \dot{X}_r &= k_3 \frac{X_{aa} X_r}{V}, & \text{(6c)} \\ V(t) &= \rho_{aa} X_{aa} + \rho_e X_e + \rho_r X_r. & \text{(6d)} \end{aligned}$$

In this reaction network, amino acids are produced by the catalytic action of the metabolic enzyme from external food molecules with rate $k_1$. Metabolic and ribosomal proteins are synthesized from the amino acids in reactions catalyzed by ribosomes with rate $k_2$ and $k_3$, respectively. It is easily verified that the model obeys the scalability condition, and therefore exhibits an exponential attractor. The volume instantaneous growth rate is computed explicitly for this system as

$$\alpha = \rho_{aa}(k_1 Y_e - (k_2 + k_3) Y_{aa} Y_r) + \rho_e k_2 Y_{aa} Y_r + \rho_r k_3 Y_{aa} Y_r. \tag{7}$$

The dynamic equations for the concentrations in this case are

$$\begin{aligned} \dot{Y}_{aa} &= k_1 Y_e - (k_2 + k_3)\, Y_{aa} Y_r - \alpha(\mathbf{Y}) Y_{aa} & \text{(8a)} \\ \dot{Y}_e &= k_2 Y_{aa} Y_r - \alpha(\mathbf{Y}) Y_e, & \text{(8b)} \\ \dot{Y}_r &= k_3 Y_{aa} Y_r - \alpha(\mathbf{Y}) Y_r, & \text{(8c)} \end{aligned}$$

with fixed point found by setting $\dot{Y}_i = 0$ as follows

$$\overline{Y_{aa}} = \frac{\overline{\alpha}}{k_3}, \tag{9}$$

$$\overline{Y_r} = \frac{\overline{\alpha}^2}{k_1 k_2 - (k_2 + k_3)\overline{\alpha}}, \tag{10}$$

$$\overline{Y_e} = \overline{Y_r} \frac{k_2}{k_3}, \tag{11}$$

and the explicit expression for iGR at the fixed point

$$\overline{\alpha} \; = \; k_1\rho_{aa}\overline{Y_e} + (k_2\rho_e + k_3\rho_r - (k_2+k_3)\rho_{aa})\overline{Y}_{aa}\overline{Y_r}. \tag{12}$$

The above equations can be combined to obtain the iGR of a cell at the fixed point as a function of model parameters,

$$\overline{\alpha} = \frac{-(k_2k_3+k_3^2+k_1k_2\rho_{aa}) + \sqrt{(k_2k_3+k_3^2+k_1k_2\rho_{aa})^2 + 4k_1k_2k_3(k_2\rho_e+k_3\rho_r-(k_2+k_3)\rho_{aa})}}{2(k_2\rho_e+k_3\rho_r-(k_2+k_3)\rho_{aa})}. \tag{13}$$

The numerator of this expression will be negative when $(k_2\rho_e + k_3\rho_r - (k_2+k_3)\rho_{aa}) < 0$, which in turn makes the denominator also negative. As a result $\overline{\alpha}$ will be positive for all positive rate parameters, showing explicitly that the attractor is of an exponentially growing nature. In the special case of $\rho_{aa} = \rho_e = \rho_r$, i.e., if we assume the equal contribution from each species to the cell size, then the steady-state iGR will be expressed in a simpler mathematical form as follows

$$\overline{\alpha} = \frac{k_1k_2k_3}{k_1k_2 + k_3(k_2+k_3)}. \tag{14}$$

Our next step is to analyze the linear stability of the fixed point $(\overline{Y}_{aa}, \overline{Y}_e, \overline{Y}_r)$ in the concentration space. This stability is determined by the eigenvalues of the Jacobian matrix evaluated at the fixed point

$$J = \begin{pmatrix} \frac{\partial f}{\partial Y_{aa}} & \frac{\partial f}{\partial Y_e} & \frac{\partial f}{\partial Y_r} \\ \frac{\partial g}{\partial Y_{aa}} & \frac{\partial g}{\partial Y_e} & \frac{\partial g}{\partial Y_r} \\ \frac{\partial h}{\partial Y_{aa}} & \frac{\partial h}{\partial Y_e} & \frac{\partial h}{\partial Y_r} \end{pmatrix} \tag{15}$$

where the functions $f, g, h$ are the right hand side of Eqs. 8 respectively. The Jacobian matrix represents the system's linear approximation near the fixed point, and its eigenvalues give information about the local dynamics. If the eigenvalues are real negative then the fixed point is stable, i.e., following perturbations the trajectories will relax back to it. The magnitude of these real parts gives a measure of how quickly this relaxation occurs, where larger magnitudes mean faster relaxation. One can obtain an exact mathematical form of the eigenvalues $\{\mathcal{E}_1, \mathcal{E}_2, \mathcal{E}_3\}$ for this reaction network as follows

$$\mathcal{E}_1 \; = \; -\overline{\alpha}, \tag{16}$$

$$\mathcal{E}_2 \; = \; -\overline{\alpha}, \tag{17}$$

$$\mathcal{E}_3 \; = \; -\left(\frac{(\overline{\alpha}^2\rho_{aa} - 2\overline{\alpha}k_3)\left[\rho_{aa}(k_2+k_3) - (k_2\rho_e + k_3\rho_r)\right] + k_3^2(k_3+k_2)}{k_3(k_2\rho_e + k_3\rho_r)}\right), \tag{18}$$

where $\overline{\alpha}$ is given in Eq. (13). The eigenvalues $\{\mathcal{E}_1, \mathcal{E}_2, \mathcal{E}_3\}$ are real negative, signifying the fixed point is stable. The absolute values of two of these eigenvalues are increasing functions of the steady-state growth rate $\overline{\alpha}$, indicating that cells with a faster growth rate will converge to the fixed point faster. A simple special case occurs if $\rho_{aa} = \rho_e = \rho_r$, i.e., if we assume equal contribution from each component to the cell size; then the third eigenvalue $\mathcal{E}_3 = -k_3$, which is real negative.

#### Supplementary Note 3: Experimental details of filamenting cells

Measurements of filamentous growth of cells were carried out using the *E. coli* K-12 derivative MG1655. Cells were transfected with two plasmids, pZA3R-mcherry - expressing a red fluorescent protein under the control of the $\lambda$-phage Pr promoter, and pRJ2001-GFP-Fis - expressing the DNA-binding protein Fis fused with GFP under the control of the lac promoter. The former plasmid was used to enable better imaging of the cell, and the latter was added to ensure that DNA was replicated during the filamentation of cells. Bacterial cultures were initiated from a frozen stock and grown overnight in LB medium at $32^0C$. The following day, the cultures were diluted in the same medium and regrown until reaching an Optical Density (OD) $\sim$0.1. They were then concentrated to an OD of 0.3 in fresh LB medium and loaded into the microfluidic device, the mother machine. Trapped cells were allowed to grow in the channels for 24 hours while supplied with LB medium flowing through the device at a rate of 1 ml/hr. Following the 24 hours of normal growth, the supplied medium was exchanged with LB containing 5$\mu$g/ml cephalexin, which induced the filamentation of the trapped bacteria. Images of the bacteria were acquired in DIC and fluorescence modes every 3 minutes during the entire experiment using a Hamamatsu ORCA-flash4.0 camera, mounted on a Nikon Eclipse Ti2 inverted microscope with a 100X objective. An Okolab microscope enclosure was used to maintain the temperature at $32^0C$. Cell length was then extracted from the acquired images using Oufti64 and custom-made MATLAB codes.

#### Supplementary Note 4: Independence of model properties from division details

The main dynamic effect described in this paper, is the disruption of concerted growth upon noisy division, and the relaxation of the multi-dimensional trajectory back to the exponential attractor. We show here that this effect is independent of details regarding the implementation of cell division in the model.

The results presented in the main text rely on a model where division is implemented as a volume threshold-crossing process. Specifically, when the cell size $V(t)$ crosses a stochastic threshold drawn from a Gaussian distribution. At division, the cell contents are distributed between the daughter cells according to a Gaussian distribution centered at a fraction $f = 1/2$ with variance $\sigma_f^2$. Fig. 1 shows three different values for this variance, extending the value chosen for display in the main text. These results clearly show that larger variance value spreads the starting point of the cell cycle further away from the fixed point, but all trajectories still converge to the same fixed point over the cell cycle.

As another extension, instead of a threshold on cell size, we considered a model where some specific protein copy number crosses a threshold to trigger division (3). We add to the model

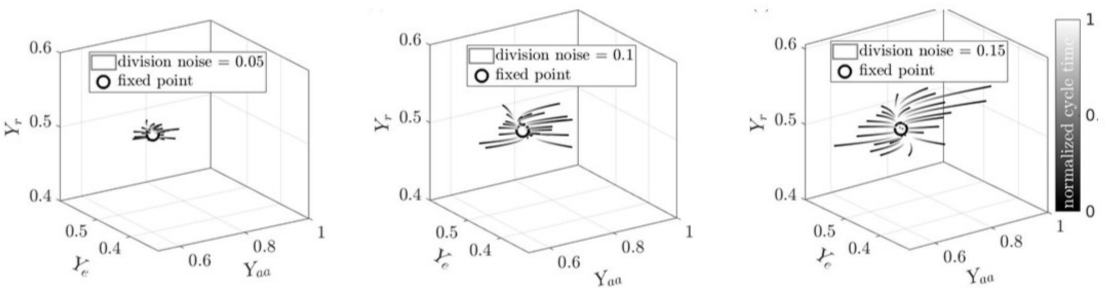


FIG. 1. **Effect of different division noise levels.** Trajectories of the dynamical system (Eqs. 6), where division is triggered by volume crossing of a stochastic threshold. At division, fractions $f_i$ and $1 - f_i$ of each components are given to each daughter cell. $f_i$ are drawn from a Gaussian distribution with three different variances as depicted in the legends. The dynamics in concentration space is shown for these three values.

equations Eqs. (6) the dynamics of this “threshold protein”, $Z$,

$$\dot{Z} = k_Z \frac{X_{aa} X_r}{V}, \tag{19}$$

neglecting the effect of this protein on the dynamics of other components (1). Also here, we see that the additional noise changes the distribution of the starting point of each cell cycle, however, the dynamics of all components along the cell cycle remains qualitatively the same (see Fig. 2).

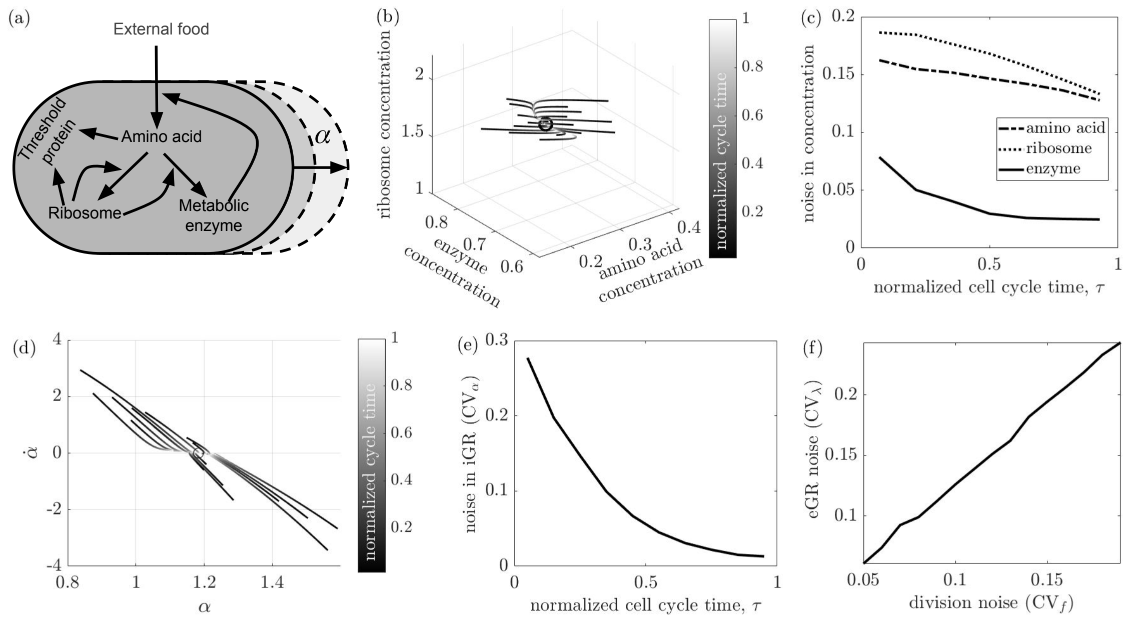


FIG. 2. **Effect of the variable triggering division - Protein number.** **(a)** Schematic of nonlinear reactions within a growing cell. Division is triggered by a threshold on the copy number of the “Threshold protein” synthesized from the amino acids with ribosomes. **(b)** Cell trajectories in the concentration phase space of the three species, with division noise of $CV_f = 0.15$ and threshold noise of 0.15. The trajectories are gray-scale coded by normalized cell cycle time from 0 (black, cycle start) to 1 (white, cycle end). **(c)** the noise in the three species concentrations as a function of cell cycle progression. **(d)** The trajectories in **(b)** depicted in the $\alpha - \dot{\alpha}$ plane. **(e)** The resulting iGR decreases over the cell cycle. **(f)** The noise in eGR increases with division noise.

### SUPPLEMENTARY FIGURES

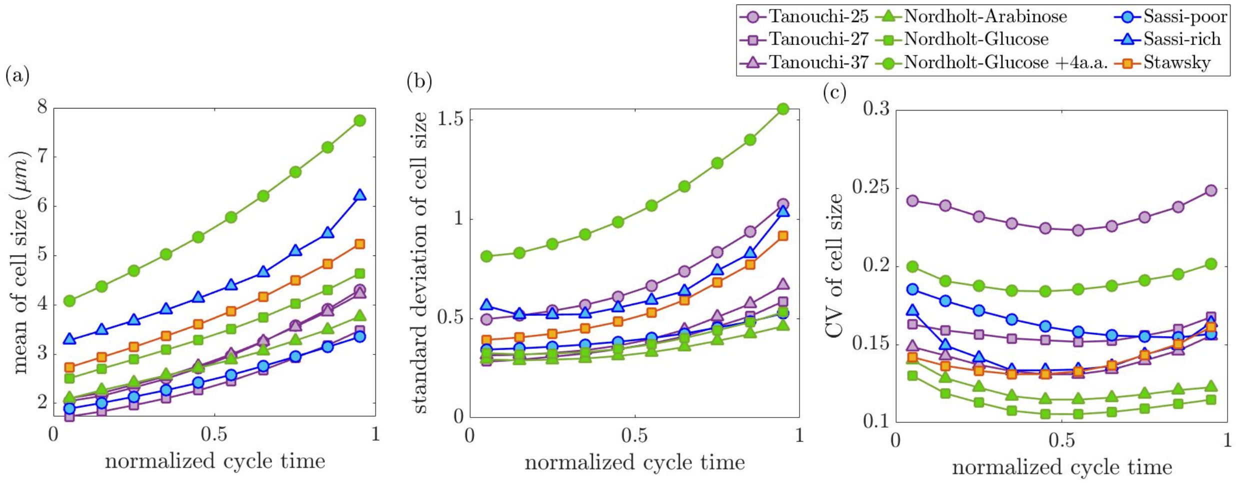


**Supplementary Figure 1:** . **Cell size statistics across the cell cycle from *E. coli* data. (a)** mean, **(b)** standard deviation, and **(c)** noise (CV) of cell size along the normalized cell cycle time obtained from *E. coli* (4–6) and *B. subtilis* data (7). Experimental details are provided in Table 2.

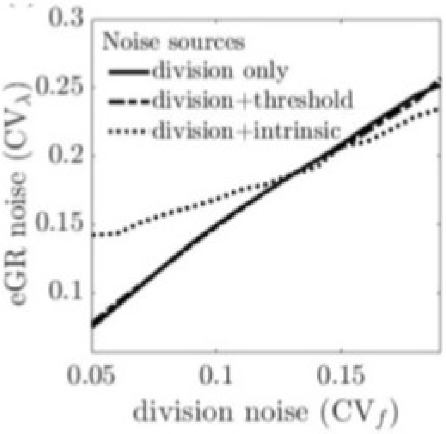


**Supplementary Figure 2:** . **Noise sources affecting growth rate variability.** $CV_\lambda$ increases monotonically with division noise $CV_f$ for fixed $\langle\lambda\rangle = 1$, which exhibits no difference for $CV_c = 0$ (solid line) and 0.15 (dashed line). This indicates that there is no effect of threshold noise on $CV_\lambda$. The effect of intrinsic noise, arising from number fluctuations, is represented by the dotted line. To incorporate this intrinsic noise we perform the Gillespie simulation (8) with exponentially distributed waiting times for each reaction in Eq. (6).

### SUPPLEMENTARY TABLES

#### Supplementary Table 1: Details of model parameters.

| **Parameters** | **Values** | **Parameters** | **Values** |
|---|---|---|---|
| $\langle V_c \rangle$: Average size threshold | 10 $\mu m$ | $\mathrm{CV}_c$: noise of size threshold | 0.15 |
| $\langle f \rangle$: Average division fraction | 0.5 | $\mathrm{CV}_f$: noise of division fraction | 0.15 |
| $k_1$: Metabolic rate of amino acid production | 4 $\mathrm{hr}^{-1}$ | $k_2$: Catalytic rate of enzyme production | 2 $\mathrm{hr}^{-1}$ |
| $k_3$: Catalytic rate of ribosome production | 4 $\mathrm{hr}^{-1}$ | $\rho_{aa}$: Molar vol. of amino acid | 1.3 |
| $\rho_r$: Molar vol. of ribosomes | 0.25 | $\rho_e$: Molar vol. of metabolic enzyme | 0.65 |

**Supplementary Table 2: Details of experimental data.**

| Strain | growth medium | #cell cycles | Temp. | mean birth size ($\mu m$) | mean growth rate ($min^{-1}$) | References |
|---|---|---|---|---|---|---|
| *E. coli* in NCM3722 background | glycerol | $1.12 \times 10^4$ | $37\ ^0C$ | 2.07 | 0.0189 | (9) |
| | sorbitol | 7439 | | 2.26 | 0.0187 | |
| | glucose | $1.25 \times 10^4$ | | 2.1 | 0.0270 | |
| | glucose + 6 a.a. | $1.68 \times 10^4$ | | 2.35 | 0.0330 | |
| | glucose + 12 a.a. | $1.33 \times 10^4$ | | 2.87 | 0.0372 | |
| | Synthetic rich | $1.09 \times 10^4$ | | 3.33 | 0.0433 | |
| | Tryptic soy broth | $1.11 \times 10^4$ | | 3.98 | 0.0569 | |
| *E. coli* in NCM3722 background | arginine | 1701 | $37\ ^0C$ | 1.48 | 0.0067 | (3) |
| | glucose + 12 a.a. | 1464 | | 2.82 | 0.0234 | |
| | glucose | 1432 | | 1.89 | 0.0144 | |
| *E. coli* in MG1655 background | M9 acetate | 1554 | | 1.88 | 0.0034 | |
| | glucose | 1807 | | 2.51 | 0.0131 | |
| | glycerol + 11 a.a. | 1491 | | 2.67 | 0.0109 | |
| *E. coli* in MC4100 background | LB minimal | 4550 | $25\ ^0C$ | 1.96 | 0.0121 | (5) |
| | | 3780 | $27\ ^0C$ | 1.64 | 0.0149 | |
| | | $1.12 \times 10^4$ | $37\ ^0C$ | 1.98 | 0.0253 | |
| *B. subtilis* | arabinose | $1.58 \times 10^4$ | $37\ ^0C$ | 2.07 | 0.0062 | (7) |
| | glucose | $1.25 \times 10^4$ | | 2.477 | 0.0108 | |
| | glucose + 4 a.a. | 2887 | | 4.105 | 0.0133 | |
| *E. coli* in MG1655 background | poor | $9.56 \times 10^4$ | $30\ ^0C$ | 1.93 | 0.0086 | (4) |
| | rich (RDM) | $4.64 \times 10^4$ | | 3.28 | 0.0153 | |
| *E. coli* in STK13 background | alanine-TrEl | 215 | $28\ ^0C$ | 1.75 | 0.0033 | (10) |
| | glucose-Cas | 409 | | 2.01 | 0.0099 | |
| | mannose | 401 | | 1.5 | 0.0035 | |
| | glucose | 344 | | 1.8 | 0.0061 | |
| | glycerol-Cas | 420 | | 1.9 | 0.0074 | |
| | glycerol-TrEl | 423 | | 1.61 | 0.0046 | |
| | glycerol | 302 | | 1.7 | 0.0042 | |
| *E. coli* in JM85 background | glucose-Cas | 406 | | 2.1 | 0.0092 | |
| | glycerol | 388 | | 2.01 | 0.0044 | |
| Wild type MG1655 *E. coli* under the control of *lac* promoter | M9 minimal | 381 | $30\ ^0C$ | 2.31 | 0.0229 | (11) |
| Wild type MG1655 *E. coli* | LB minimal | 4096 | $30\ ^0C$ | 2.65 | 0.0357 | (12) |
| Wild type MG1655 *E. coli* under the control of $\lambda$ promoter | M9 minimal | 1794 | | 1.35 | 0.0509 | |
| | LB minimal | 3551 | | 2.78 | 0.0252 | |
| Wild type MG1655 *E. coli* under the control of *lac* promoter | M9 minimal | 562 | | 2.43 | 0.0233 | |
| Wild type MG1655 *E. coli* under the control of both $\lambda$ & *lac* promoter | LB minimal | 188 | | 1.21 | 0.0257 | |
